# S100A4 plays a role in mouse arterial smooth muscle cell motility. Implication for intimal thickening formation

**DOI:** 10.1101/2022.09.30.510253

**Authors:** LM. Cardoso dos Santos, J. Klingelhöfer, ML. Bochaton-Piallat

## Abstract

During atherosclerosis, smooth muscle cells (SMCs) migrate and accumulate in the intima, where they switch from a contractile to a synthetic phenotype. This process is associated with decreased expression or loss of contractile proteins, such as α-smooth muscle actin (α-SMA) or smooth muscle myosin heavy chains (SMMHCs). We previously demonstrated that S100A4, a small calcium-binding protein, which exhibits intra- and extracellular functions, is a marker of the synthetic SMC phenotype. We have recently shown that neutralization of extracellular S100A4 in an ApoE knockout (KO) mouse model with established atherosclerotic lesions decreased the overall atherosclerotic burden. To explore the role of S100A4 in the accumulation of SMCs in the intima, we induced intimal thickening (IT) formation in full S100A4 knockout (KO) and wild type (WT) mice by completely ligating the left common carotid artery. With this model, we generated SMC-rich lesions. The deletion of S100A4 did not influence the size of the IT, neither its composition, as assessed by the expression of α-SMA and SMMHCs in S100A4 KO animals compared with WT animals, 4 weeks after ligation. Using primary cells isolated from both strains, we demonstrated that S100A4 KO SMCs were less prone to migrate than WT SMCs but they did not differ in their proliferative capacity. Our results indicate that S100A4 plays a role in SMC motility *in vitro* but its deletion does not influence IT formation in the mouse carotid artery ligation model.

## Introduction

Atherosclerosis is the main cause of cardiovascular disease. During the development of the lesion, a gradual accumulation of lipids and inflammatory cells takes place in the intima of arteries. Moreover, smooth muscle cells (SMCs) migrate from the media to the intima, proliferate and undergo a transition from contractile to synthetic phenotype.^1^ This process is characterized by the dedifferentiation of SMCs, which partially lose their contractile machinery (i.e. downregulation or complete loss of cytoskeletal protein, such as α-smooth muscle actin [α-SMA], smooth muscle myosin heavy chains [SMMHCs], and transgelin [also known as SM22α]).^2^ During the last two decades, SMCs were considered as beneficial players since it was thought that their main function was to protect the plaque from rupture, by contributing to the formation of the fibrous cap.^2^ This notion shifted with the understanding that SMC switching is not a binary process and that, due to their immense plasticity, SMCs exhibit a spectrum of phenotypes, including a macrophage-like phenotype.

From normal porcine coronary arteries, our group isolated two distinct SMC populations, spindle-shaped (S) and rhomboid (R), corresponding to the contractile and synthetic phenotypes, respectively.^3^ The R-SMC population exhibits greater proliferative and migratory capacity, compared with S-SMCs. Moreover, S-SMCs, in response to platelet-derived growth factor (PDGF)-BB or fibroblast growth factor 2, switched towards a R-phenotype, in a reversible way.^3^ Later on, using a 2D gel electrophoresis approach, we identified S100A4 as specifically expressed in R-SMCs but not in S-SMCs.^4^ We and others further demonstrated that S100A4 is expressed in intimal SMCs *in vivo* in humans, pigs, and mice.^4-9^

S100A4 is a small calcium-binding protein belonging to the S100 family. It exists in a variety of conformations, oligomers being the most active extracellular form.^10,11^ It exhibits intra- and extracellular functions, and is expressed and released by a variety of cells, such as cancer cells, fibroblasts and immune cells (e.g. macrophages).^11^ S100A4’s strongest association is with cancer and metastasis; however, it has also been described in the context of fibrosis, cardiac hypertrophy, and rheumatoid arthritis.^12,13^ Inside the cell, it interacts with many players, mainly the ones involved in cytoskeletal organization and motility.^14,15^ In addition, S100A4 interacts directly with the oncogene p53, interfering with its DNA binding capacity.^16^ Noteworthy, the deletion of S100A4 in murine cancer models greatly decreased the metastatic potential.^11^ Extracellularly, S100A4 acts through different receptors, mainly related to inflammatory pathways, such as the receptor for advanced glycation end-products (RAGE) or the toll-like receptor 4 (TLR4), inducing activation of the nuclear factor kappa B (NFκB), and production of matrix metalloproteinases (MMPs) as well as pro-inflammatory proteins.^17-19^ We have recently shown that oligomeric S100A4 activates NFκB through TLR4 and should be co-stimulated with PDGF-BB for a complete transition of porcine coronary artery SMCs towards a pro-inflammatory profile.^6^ Moreover, *in vivo*, using a neutralizing antibody against S100A4,^20^ we were able to decrease the size of the lesions and the systemic and local inflammation. S100A4 neutralization also led to stabilization of atherosclerotic plaques, through an increase in α- SMA-positive intimal SMCs.^6^

In the current study, we aimed at exploring the role of S100A4 in the accumulation of SMCs during the development of intimal thickening (IT). For this purpose, we used full S100A knockout (KO) mice and wild type (WT) mice as controls,^21^ where we induced IT after carotid artery ligation. Four weeks after ligation, we observe that S100A4 deletion does not influence IT formation. *In vitro*, using mouse primary SMCs, we show that the migratory activity decreases in S100A4 KO SMCs when compared with WT SMCs.

## Methods

### Cell culture, treatments and reagents

Aortic SMCs were isolated from 5 to 8 weeks-old WT and S100A4 KO mice. Mice were euthanized by CO_2_ asphyxiation and perfused (intracardiac) with 0.9% NaCl. Aortas were collected until the inguinal bifurcation and placed on ice, in isolation medium (i.e. DMEM [#61965-026, Gibco, Life Technologies], 2% HEPES [#15630-056, Gibco, Life Technologies], 100 U/ml penicillin and 100 μg/ml streptomycin [#150070-063 Gibco, Life Technologies]). Aortas were washed 3 times in isolation medium, cleaned of all fat and digested for 10 min at 37°C in digestion medium (i.e. isolation medium supplemented with 1.5 mg/mL of collagenase type 2 [#LS004176, Worthington-Biochem] and 0.7 U/mL of elastase type III [#E0127, Sigma-Aldrich]). The adventitia was then mechanically removed, and the aorta cut longitudinally to remove blood clots and endothelial cell (EC) layer, followed by washing in isolation medium 3 times. Aortas were cut in small pieces (1.5 to 3 mm) and further incubated in digestion medium at 37°C with gentle rotation for 20 min, triturated 15 times, and allowed to digest for another 20 min. Digestion was stopped by adding isolation medium supplemented with 10% fetal calf serum (FCS, 2-01F10-I, Bioconcept). The cell solution was triturated 15 times and centrifuged at 600g for 5 min. The pellet was then resuspended in SMC medium (i.e. DMEM/Nutrient Mixture F-12 GlutaMAX™ [#31331-028, Gibco, Life Technologies], 100 U/ml penicillin and 100 μg/ml streptomycin, supplemented with 10% FCS), plated in 6 cm petri dishes (4 aortas/dish) for approximately 7 days. Cells were passed at a 1:2 ratio when 80% to 90% confluency was reached. Cells were cultured at 37°C 5% CO_2_, passed at a 1:2 split ratio, and used between passages 3 and 8.

For the treatments, cells were plated for 24 h in SMC medium supplemented or not with FCS, before adding the different compounds. Murine PDGF-BB (#315-18, PeproTech Nordic) was used at a concentration of 20 ng/mL. Human recombinant N-terminal His-Tag S100A4 (rS100A4; kindly provided by Dr Jörg Klingelhöfer, University of Copenhagen) was produced in a bacterial expression system and purified by affinity chromatography. Different fractions of S100A4 conformations were isolated by size exclusion chromatography. The fractions corresponding to molecular masses ranging from 44 to 200 kDa were pooled and considered as the active form of S100A4.^22^ rS100A4 was tested at 1, 2.5 and 5 μg/mL, with the concentration of 2.5 μg/mL chosen for all experiments.

### Animals and carotid artery ligation

All animal studies were performed after approval by the Swiss Federal Veterinary Office and were in accordance with the established Swiss guidelines and regulations. These animal studies conform to the guidelines from Directive 2010/63/EU of the European Parliament on the protection of animals used for scientific purposes. For animals that were subjected to a surgical intervention, their welfare was assessed daily during the first week, by weighting and observing animal’s behavioral features, and weekly for the following period of time.

Twenty-six week-old male WT (n=12) and S100A4 KO (n=10) mice^21^ were subjected to left carotid artery ligation, a procedure adapted from Kumar and Lindner^23^. Briefly, animals received preoperative analgesia with subcutaneous injection of buprenorphine (0.1 mg/kg body weight) and were then anesthetized using 2 to 4% isofluorane by inhalation. Using a stereoscope equipped with a thermal pad to avoid hypothermia, an incision was made in the neck region, and the left carotid artery was exposed and completely ligated under the bifurcation with a silk 6-0 non-absorbable suture (#65201F, Assut Sutures). Post-operative analgesia with subcutaneous injection of buprenorphine (0.1 mg/kg body weight) was maintained for 2 to 3 days after surgery. All animals received aspirin via drinking water (16mg/kg/day) throughout the entire duration of the experiment. Black bottles were used and the water was changed every 2 to 3 days, to avoid aspirin degradation. Mice were fed with normal chow diet and sacrificed 4 weeks after surgery following CO_2_ asphyxiation. After being perfused with 0.9% NaCl, ipsi- and contralateral carotid arteries were dissected, cleaned of the fat by using a Stemi 508 stereoscope (Carl Zeiss), and snap-frozen in OCT compound (#361603E, VWR Chemicals). For histological (haematoxilin/eosin and Prussian blue staining) and immunofluorescence purposes, carotid arteries were completely cryosectioned with 5 μm-thick sequential sections starting between 50 to 100 μm from the ligature point until the end of the carotid artery, and placed on SuperFrost Plus™ slides.

### Immunofluorescence

Immunofluorescence staining was performed on adherent mouse SMC cultures and carotid artery cryosections. Primary antibodies against α-SMA (mouse monoclonal IgG2a, 1:50, clone 1A4)^24^, SMMHCs (rabbit polyclonal IgG, 1:50, BT-562, Biomedical Technologies), S100A4 (rabbit polyclonal IgG, 1:200, A5114, DAKO), and CD68 (rat monoclonal IgG2a, 1:5000, FA-11, Abcam) were used. Cells and cryosections were fixed for 15 min in 1% paraformaldehyde (PFA), then rinsed in phosphate buffer saline (PBS), and further incubated for 5 min in ice-cold methanol, except for anti-CD68, for which cryosections were fixed in ice-cold acetone for 5 min. After washing in PBS, cells and cryosections were stained with the primary antibody for 1h at room temperature (RT) and rinsed in PBS. The following secondary antibodies were used: Alexa Fluor® 488-conjugated goat anti-mouse IgG2a, Alexa Fluor® 488- or 594-conjugated goat anti-rabbit IgG (Jackson ImmunoResearch); and fluorescein-conjugated rabbit anti-rat IgG (Vector Laboratories) and applied for 1 h at RT, together with 4⍰,6-diamidino-2-phenylindole (DAPI, Fluka). Slides were mounted in buffered polyvinyl alcohol (PVA) and images were taken using a fluorescence microscope (Axioskop 2, Carl Zeiss) equipped with a x20/1.4 objective and a high sensitivity, high-resolution digital color camera (Axiocam 702, Carl Zeiss). The image analysis was done using Zen 3.3 blue edition (Carl Zeiss) and ImageJ v1.53c software.

For NFκB activation, 100’000 cells were plated in SMC medium (n=3) and allowed to adhere for 24 h; then rS100A4 was added to the medium at 1, 2.5 and 5 μg/mL. After 1 h, cells were fixed for 15 min in 1% PFA, rinsed in PBS and stained with mouse monoclonal NFκB p65 IgG1 (1:50, sc8008, Santa Cruz Biotechnology) followed by Alexa Fluor® 488-conjugated goat anti mouse IgG1 (Jackson ImmunoResearch). Lipopolysaccharides (LPS) was used as a positive control (1 μg/mL for 1h).

### Proliferation assay

For the basal proliferation assays, 140’000 cells were plated in a 6 cm petri dish for 2, 4 and 7 days in SMC medium. At the different timepoints, cells were trypsinized and counted manually using a Neubauer’s chamber (performed in duplicate with a minimum of n=4). For the treatments with PDGF-BB, rS100A4 and PDGF-BB/rS100A4, 70’000 cells were plated in a 6 cm petri dish and after 24 h, compounds were added for 4 days (n=7 and n=8, for WT and S100A4 KO SMCs, respectively).

### Migration assay

For the migration assay, we used a modified Boyden Chamber where 400’000 cells were plated in cell culture translucent inserts for 6-well plates, 8 μm-pore size, polyethylene terephthalate membranes (#657638, ThinCerts™, Greiner Bio-one), in a total of 3 mL serum-free SMC medium, and allowed to adhere for 4h. Then, either 10% FCS (performed in duplicate with a minimum of n=3), PDGF-BB, rS100A4 or PDGF-BB/rS100A4 (at least n=7) were added to the lower compartment for 18h. Cells were washed in DMEM/2% HEPES, fixed in DMEM/1% PFA for 20 min at RT, permeabilized in ice-cold methanol for 5 min, and stained with DAPI for 15 min at RT. Cells were then removed from the upper layer of the membrane using a cotton swab, the membrane detached and mounted on a glass slide using PVA. Membranes images were taken on a fluorescence microscope (Axioskop 2) equipped with a x20/1.4 objective and a high sensitivity, high-resolution digital color camera (Axiocam 702). Six random fields were acquired from each membrane and the number of cells was quantified manually using ImageJ v1.53c software. Results are expressed as an average number of cells per field.

### Protein extraction, electrophoresis and Western blotting

Cells were trypsinized and proteins were extracted as previously described.^4^ Briefly, cells were centrifuged at 600g for 5 min, washed in serum free SMC medium complemented with protease inhibitors (#4693132001, Roche) and centrifuged at 600g for 5 min. Pellets were lysed in sample buffer (i.e. 80 mM Tris-HCl pH 6.8, 0.1 M dithiothreitol, 2% sodium dodecyl sulfate [SDS], 10% glycerol, 0.001% bromophenol blue), and boiled for 5 min. Protein concentration of SMC lysates was determined using the BCA assay (Thermofisher), according to manufacturer’s instructions.

For Western blotting, 1 or 10 μg of total protein were loaded and separated by SDS–PAGE on 10% mini gels (Bio-Rad), and transferred into a 0.2 μm-pore size nitrocellulose membrane (#GE10600001, Amersham™ Protran®, GE Healthcare). Membranes were then quickly incubated in Ponceau Red to confirm proper protein migration, and blocked with PBS 5% non-fat dried milk for 1 h at RT. Primary antibodies against α-SMA (1:500), SMMHCs (1:1000), and S100A4 (1:400) were incubated in PBS/0.1% Triton X-100 (PBS-T)/3% BSA for 1 h at RT, washed 3 times in PBS-T. Horseradish peroxidase-conjugated goat anti-mouse or anti-rabbit IgG were used as secondary antibodies and incubated for 1h RT in PBS-T/0.3% BSA. After washing 3 times in PBS-T, WesternBright Sirius HRP substrate (#K-12043-D10, Advansta) was used for chemiluminescence detection. Signals were digitized with a LAS4000 mini luminescent image analyzer (Fujifilm) and analyzed by using ImageJ v1.53c software. Results were normalized to GAPDH expression.

### Statistical analysis

Statistics were performed in Graph Pad Prism 9 by using the two-tailed or multiple unpaired Student’s t-tests with a confidence level of 95%. Multiple group comparisons were performed by using one-way ANOVA, followed by Tukey’s multiple comparison test. Results are shown as mean ± 95% confidence internal (CI). Differences were considered statistically significant at values of *p≤0.05.*

## Results

### Morphology and SMC differentiation marker characterization

Phase-contrast analysis showed that WT and S100A4 KO SMCs exhibited similar heterogeneous morphology with diverse cell size and shape **(Figure 1A).** Immunofluorescence staining for α-SMA and SMMHCs showed that both SMC populations were well differentiated and displayed α-SMA-positive stress fibers **(Figure 1B).** Approximately 10 to 15% of cells also exhibited SMMCHs typical punctuated distribution **(Figure 1B,** right panel), a marker of mature SMCs. As expected, S100A4 was absent in S100A4 KO SMCs and slightly expressed in WT SMCs **(Figure 2A,** control conditions).

**Figure 1.**
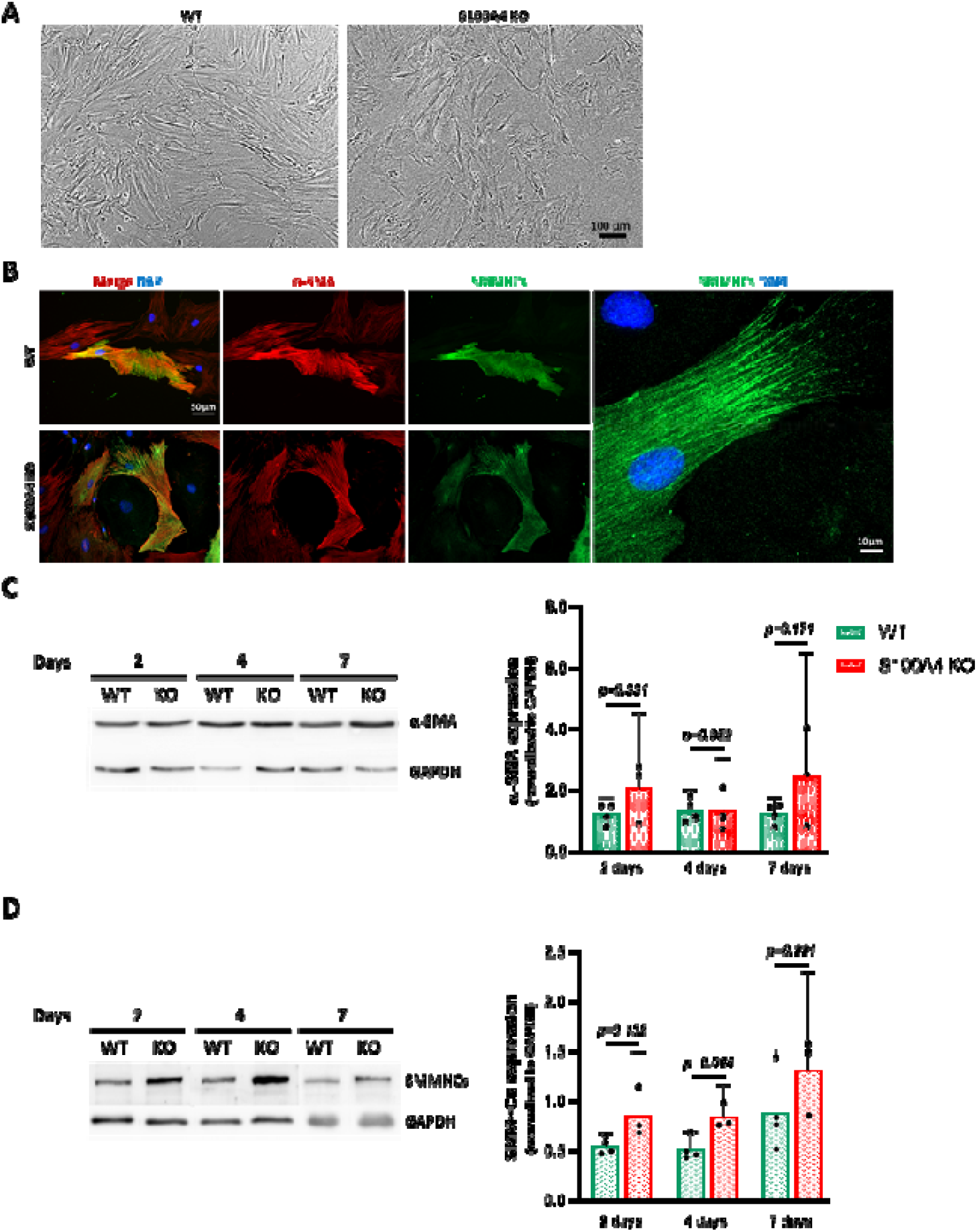
Morphological features and SMC differentiation marker expression in WT and S100A4 KO SMC populations. **(A)** Phase-contrast photomicrographs of WT and S100A4 KO SMCs. **(B)** Double-immunofluorescence staining for α-SMA and SMMHCs for WT and S100A4 KO SMCs, depicting the organization of stress fibers and the punctuated pattern of SMMHCs (higher magnification, right panel). Representative Western blots from WT and S100A4 KO SMCs, and respective quantification of **(C)** α-SMA and **(D)** SMMHCs in WT and S100A4 KO SMCs after 2, 4 and 7 days in culture. In **(B)** nuclei are stained in blue by DAPI. In **(A)** and **(B)** n=12 and 10 for WT and S100A4 KO SMCs, respectively. In **(C)** and **(D)** n=4 and 3 for WT and S100A4 KO SMCs, respectively.

**Figure 2.**
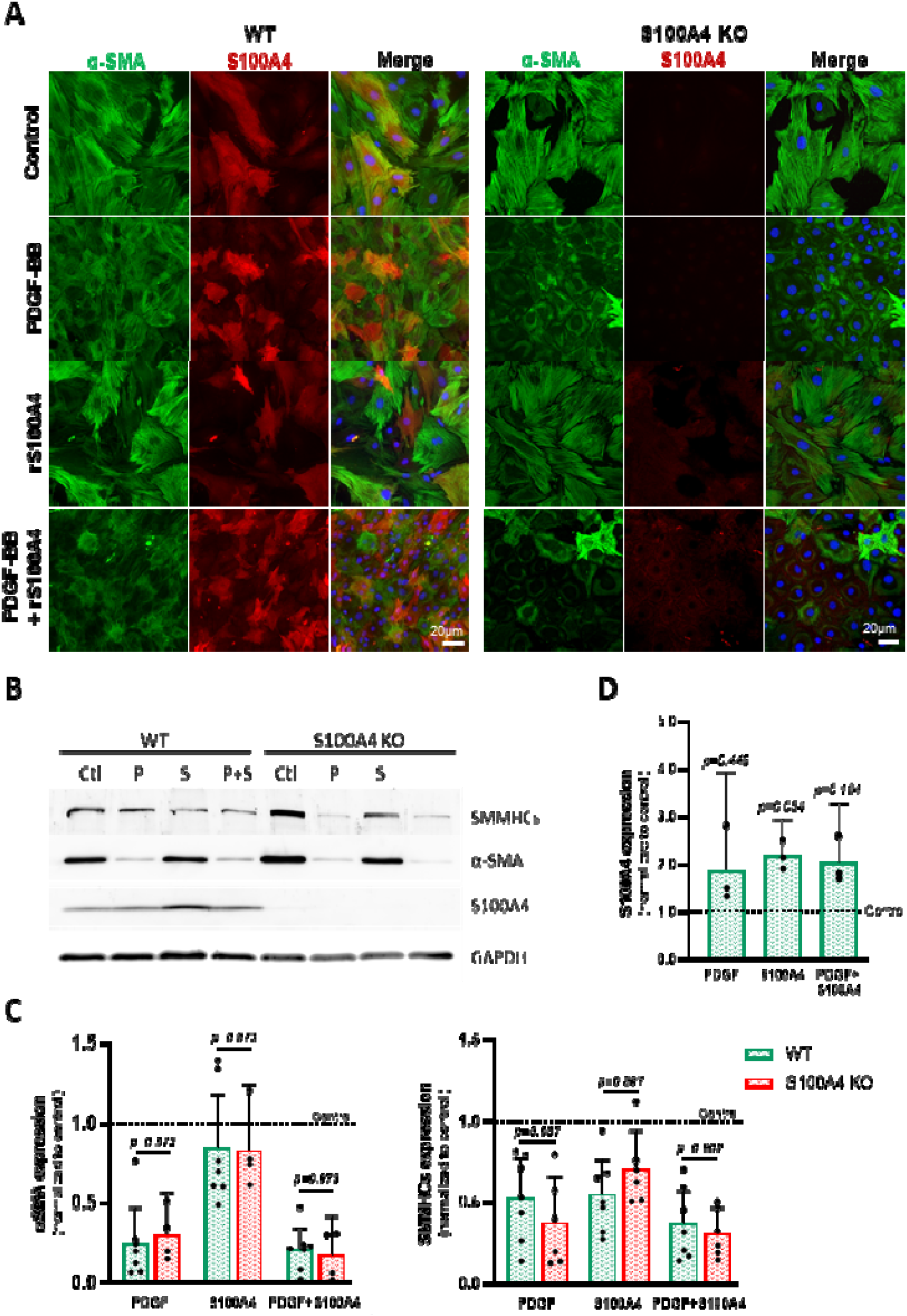
Effect of PDGF-BB, rS100A4 or PDGF-BB/rS100A4 on SMC differentiation marker expression in WT and S100A4 KO SMCs. **(A)** Double-immunofluorescence staining for α-SMA and S100A4 showing WT and S100A4 KO SMCs treated with PDGF-BB, rS100A4 or PDGF-BB/rS100A4 for 4 days. **(B)** Representative Western blots for α-SMA, SMMHCs and S100A4, and **(C)** corresponding quantification, relative to control (dashed line), of α-SMA, SMMHCs and **(D)** S100A4 in WT and S100A4 KO SMCs. S100A4 was not detected in S100A4 KO SMCs, as expected. In **(A)** nuclei are stained in blue by DAPI. In **(C)** n=7 and n=4 for α-SMA for WT and S100A4 KO SMCs, respectively, and n=7 for SMMHCs for both SMC populations. In **(D)** n=3 for both SMC populations.

Western blots showed no significant difference in α-SMA and SMMHCs expression at 2, 4 and 7 days of culture for WT as well as S100A4 KO SMCs **(Figure 1C** and D). However, we observed a trend for higher expression of SMMHCs in S100A4 KO SMCs compared with WT SMCs, in particular at 2 and 4 days in culture. S100A4 was absent in KO SMCs while it was present at low-level in WT cells **(Figure 2B,** control conditions), confirming immunofluorescence staining results. Therefore, WT and S100A4 KO SMCs do exhibit similar level of differentiation.

PDGF-BB is a known promoter of SMC phenotypic transition from a contractile towards a synthetic phenotype. In porcine coronary artery SMCs, we previously demonstrated that the combination of PDGF-BB and rS100A4 was necessary for the complete transition towards a proinflammatory phenotype model.^6^ WT and S100A4 KO SMCs were treated with PDGF-BB, rS100A4 and PDGF-BB/rS100A4 for 4 days **(Figure 2).** By immunofluorescence staining **(Figure 2A),** PDGF-BB, and to a greater extent PDGF-BB/rS100A4, induced a reduction in size and a more cobblestone morphology in the majority of the SMCs, which was associated with decreased expression of α-SMA, in both WT and S100A4 KO SMCs. In contrast, rS100A4 alone did not change cell morphology in both SMC populations. By Western blotting, we confirmed that, compared with control, PDGF-BB markedly decreased α-SMA expression, and to a lesser extent, SMMHCs expression in WT and S100A4 KO SMC populations (p=0.007 and p=0.0104 for α-SMA and p=0.0128 and p=0.0093 for SMMHCs, respectively; **Figure 2B** and **C**). PDGF-BB/rS100A4 decreased, to the same extent as PDGF-BB alone, both α-SMA and SMMHCs expression in WT and S100A4 KO SMCs (p<0.00001 and p=0.0053 for α-SMA and p=0.0012 and p=0.0004 for SMMHCs, respectively; **Figure 2B** and **C**). In contrast, rS100A4 did not affect α-SMA expression in WT and S100A4 KO SMCs and SMMHCs expression in S100A4 KO SMCs; it significantly decreased SMMCHs expression only in WT SMCs (p=0.0086, compared with control; **Figure 2B** and **C**). No difference was observed between WT and S100A4 KO SMCs when treated with PDGF-BB, rS100A4 and PDGF-BB/rS100A4 **(Figure 2B** and **C**). The expression of intracellular S100A4 was not changed by the different treatments **(Figure 2D).** Therefore, WT SMCs do not differ from S100A4 KO SMCs with respect to SMC differentiation marker expression in response to growth factors.

The NFκB pathway was previously shown to be activated by rS100A4.^6^ Both SMC populations treated with 1, 2.5 and 5 μg/mL for 1h exhibited activation of NFκB, associated with its nuclear localization **(Figure 3).** The optimal concentration was 2.5 μg/mL, the lowest concentration for which an almost complete NFκB activation was obtained in both conditions.

**Figure 3.**
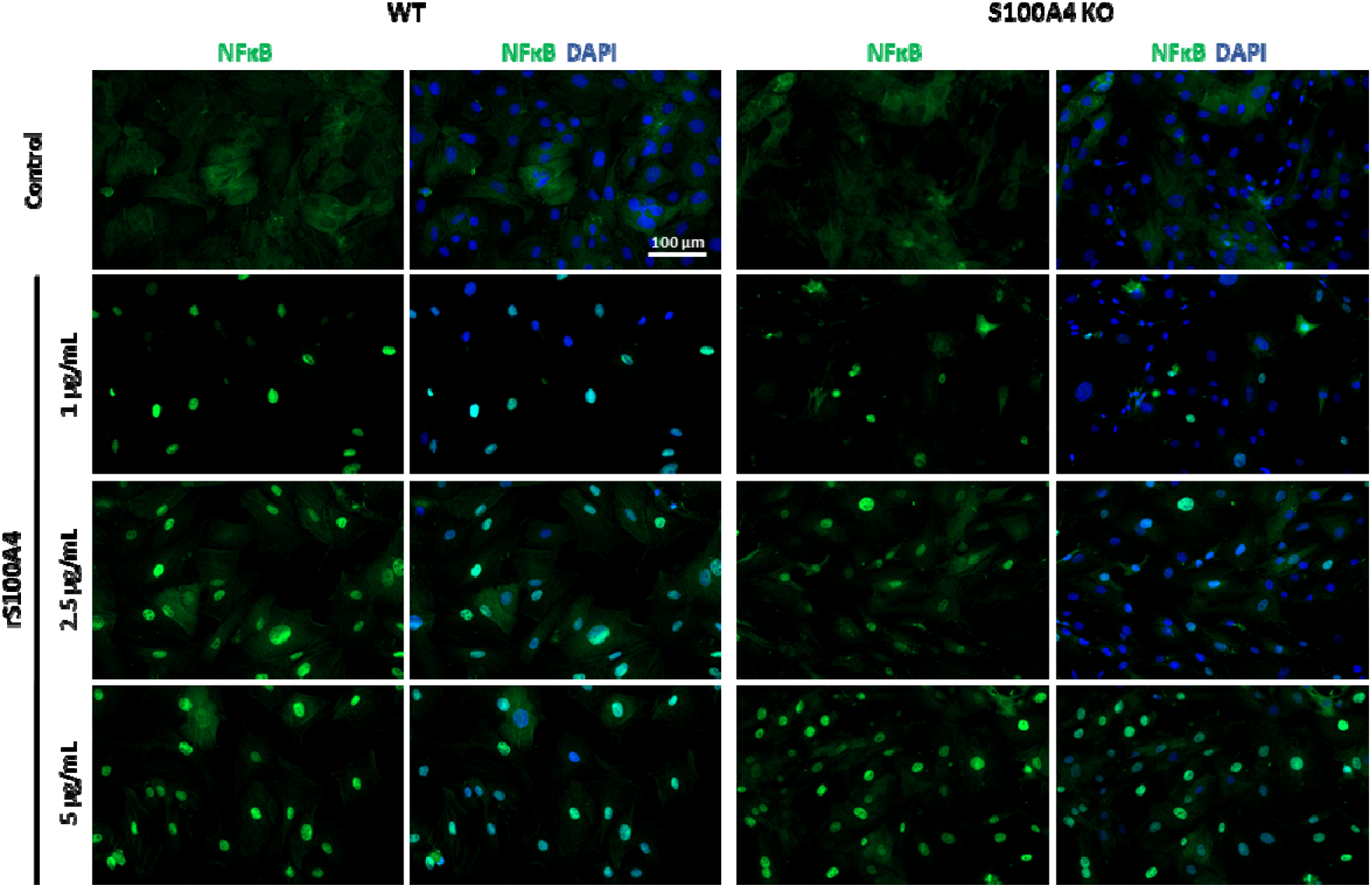
Effect of extracellular S100A4 in NFκB activation in WT and S100A4 KO SMCs. Immunofluorescence staining for NFκB showing its nuclear localization in both WT and S100A4 KO cells treated with 1, 2.5 and 5 μg/mL of rS100A4. Nuclei are stained in blue by DAPI. n=3 for both SMC populations.

### Cell proliferation

The number of WT and S100A4 KO SMCs progressively and significantly increased over time, in particular from 4 to 7 days (p=0.0366 and p=0.0125 for WT and S100A4 KO SMCs, respectively). However, no difference in proliferative capacity was observed between them on basal conditions at 2, 4 or 7 days **(Figure 4A).** PDGF-BB or PDGF-BB/S100A4, but not rS100A4 alone, similarly increased cell proliferation in both SMC populations compared with control **(Figure 4B).** Therefore, WT SMCs do not differ from the S100A4 KO SMCs with respect to their basal proliferation or ability to proliferate in response to growth factors.

**Figure 4.**
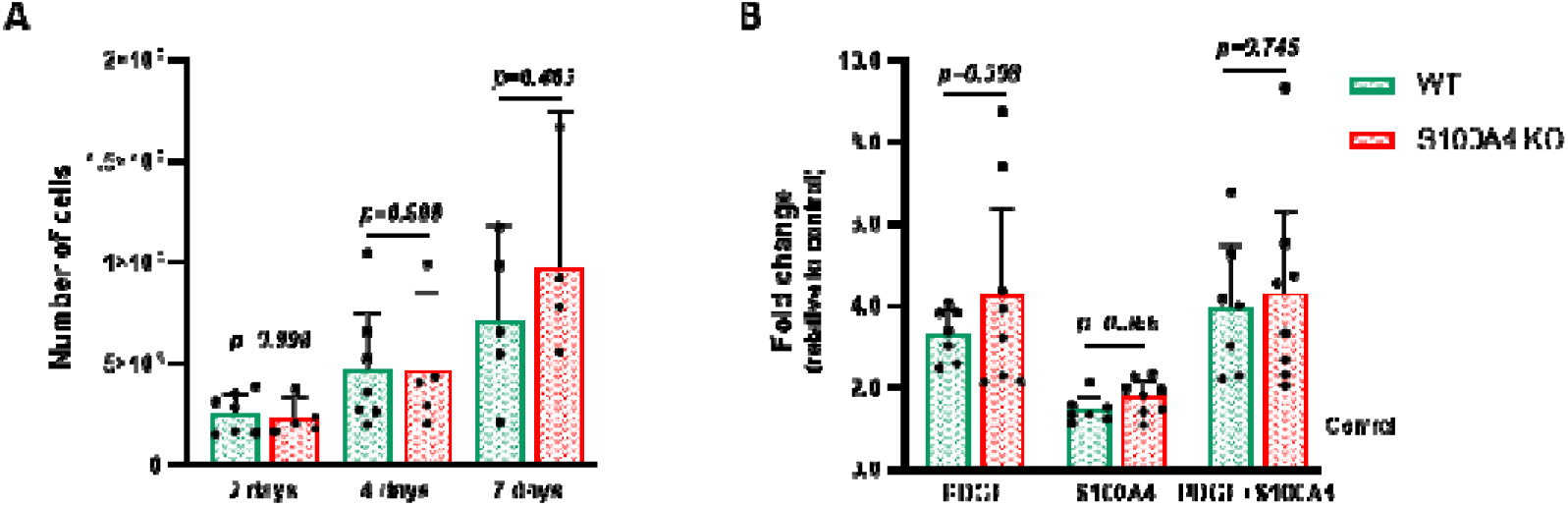
Proliferative capacity of WT and S100A4 KO SMCs. Graphical representation of the absolute number of cells in WT and S100A4 KO SMCs, after (A) 2, 4, and 7 days in culture. (B) Graphical representation of the fold change, relative to control (dashed line), of the number of WT and S100A4 KO SMCs upon treatment with PDGF-BB, rS100A4 or PDGF-BB/rS100A4. In (A) n=5 and n=4, and in (B) n=7 and n=8 for WT and S100A4 KO, respectively.

### Cell migration

By using a Boyden’s chamber, we counted the number of cells that completely migrated from the upper side of the membrane. S100A4 KO SMCs exhibited decreased migratory capacity compared with WT SMCs when cultured in 10% FCS **(Figure 5A).** As expected, PDGF-BB significantly increase migration in WT SMCs, when compared with serum-free medium control condition (p=0.003) and rS100A4 (p=0.009). PDGF-BB treatment in S100A4 KO SMCs only significantly differed from that of control (p=0.003). The combination of PDGF-BB/rS100A4 increased migration to a greater extent, being significantly different from control (p=0.001 and p=0.003) and rS100A4 (p=0.003 and p=0.005), in WT and S100A4 SMC, respectively **(Figure 5B).** Nonetheless, no difference was observed between PDGF-BB and PDGF-BB/rS100A4 **(Figure 5B).** As for rS100A4 alone, it did not change cell migration when compared with control. Interestingly, in all conditions studied, the migratory activity of S100A4 KO SMCs was markedly decreased than that of WT SMCs **(Figure 5B).** Therefore, the deletion of S100A4 in SMCs led to a lower migratory capacity, in both basal conditions or upon treatments with PDGF-BB and/or rS100A4.

**Figure 5.**
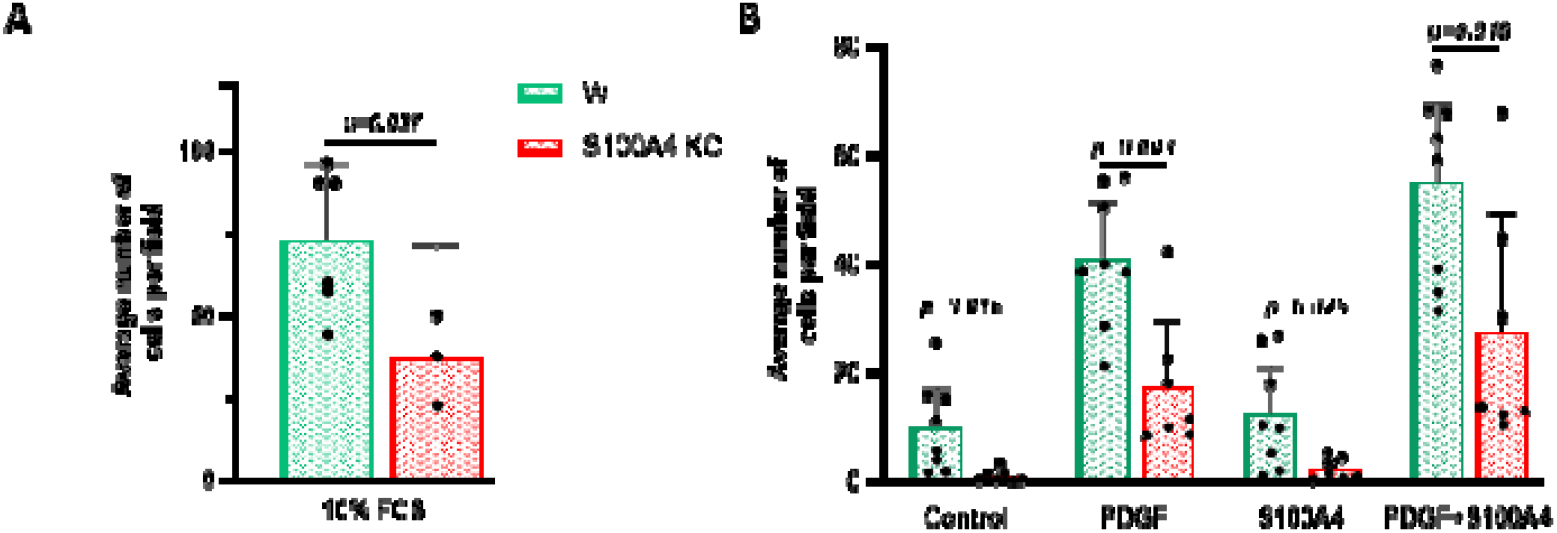
Migratory capacity of WT and S100A4 KO SMCs. Graphical representation of the number of migrating SMCs in WT and S100A4 KO SMCs treated with **(A)** 10% FCS, **(B)** serum-free medium (control), PDGF-BB, rS100A4 or PDGF-BB/rS100A4. In **(A)** n=6 and n=3, and in **(B)** n=7 and 9 for WT and S100A4 KO SMCs, respectively.

### Carotid artery ligation

To investigate whether the deletion of S100A4 affects the accumulation of SMCs in the intima, left common carotid artery ligation was performed in male WT (n=12) and S100A4 KO (n=10) mice at 26 weeks of age, for 4 weeks. Ipsilateral and contralateral carotid arteries were collected and analyzed. Neither IT area, media area or media thickness were significantly different in S100A4 KO mice compared with WT mice **(Figure 6A and B).** In a small percentage of the animals, the carotid artery was almost completely occluded, with a small lumen visible in the center of the lesion. In such cases, some areas of the lesions were devoid of nuclei; Prussian blue staining (affinity for iron) was negative indicating the absence of a thrombus (data not shown).

**Figure 6.**
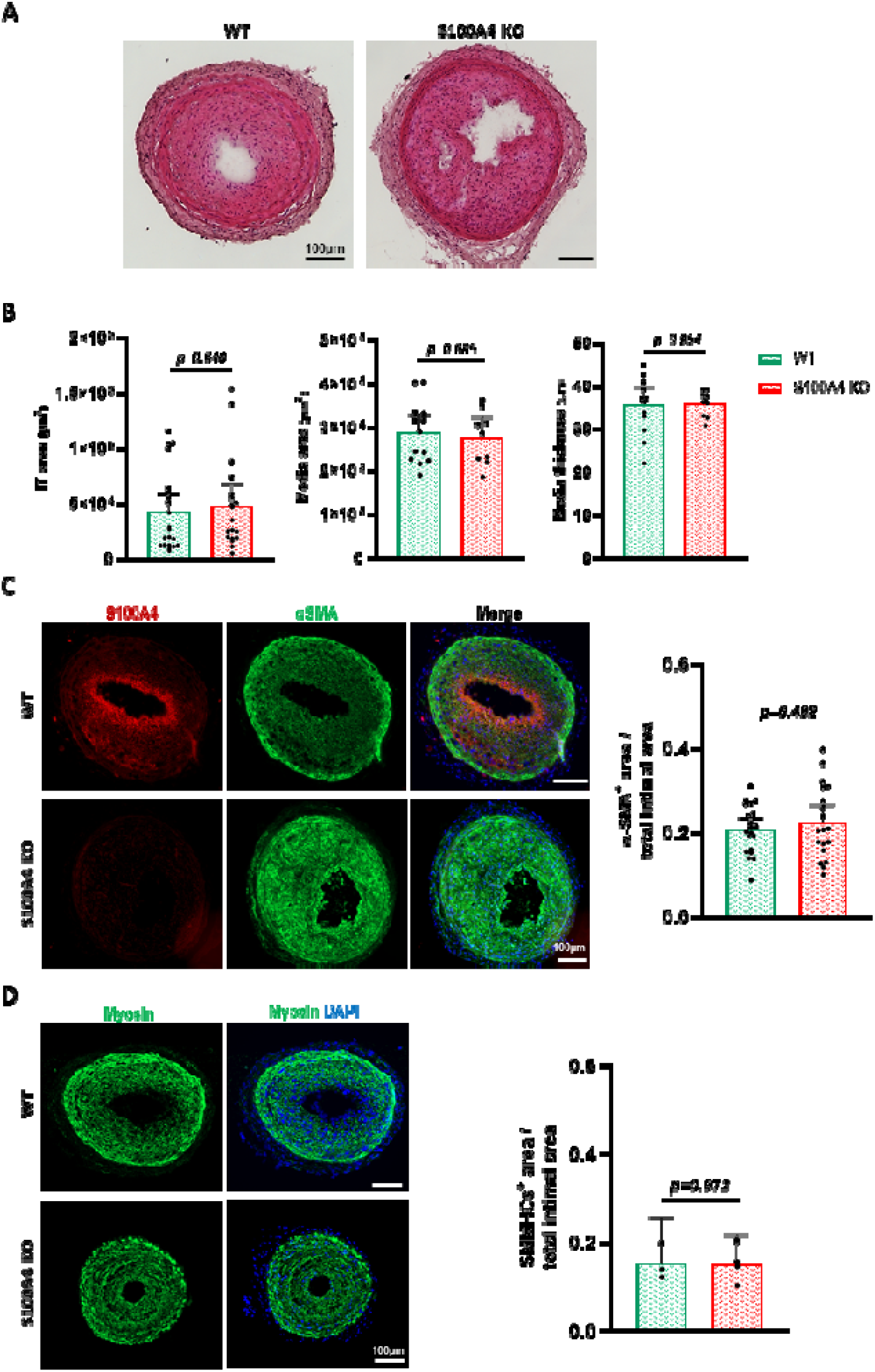
Effect of S100A4 deletion in IT formation after mouse carotid artery ligation. **(A)** Haematoxylin and eosin staining. **(B)** Quantification of the IT area, media and media thickness. Double-immunofluorescence staining for **(C)** α-SMA and S100A4 in WT and S100A4 KO animals, and α-SMA quantification. **(D)** Immunofluorescence staining for SMMHCs in WT and S100A4 KO animals, and their respective quantifications, after 4 weeks of carotid ligation. In **(C)** and **(D)**, DAPI stains the nuclei in blue. n=12 and n=11 for WT and S100A4 KO animals, respectively.

The composition of the IT, assessed by the staining with α-SMA and SMMHCs **(Figure 6C** and **D**) did not show any differences, the majority of the cells being positive for α-SMA **(Figure 6C)** and SMMHCs **(Figure 6D).** In WT animals, S100A4 staining was absent from the media layer and present in intimal cells, confirming that S100A4 is a marker of intimal cells. Interestingly, the S100A4 staining was restricted to the most luminal intimal cell layers and was prominently co-stained with α-SMA **(Figure 6C).** As expected, in S100A4 KO animals, the S100A4 staining was absent **(Figure 6D).** Overall, our results indicate that the deletion of S100A4 does not influence the IT development and progression.

## Discussion

In this study, we used full S100A4 KO and WT mice, in which we induced SMC-rich IT by completely ligating the left common carotid artery. We unexpectedly observed that, in this model, the absence of S100A4 did not impact the development of IT, 4 weeks after ligation. *In vitro*, SMCs cultured from aortas of S100A4 KO mice exhibit impaired migratory capacity compared with WT mice.

The carotid artery ligation is considered as a model of non-inflammatory restenosis, characterized by an extensive IT mostly composed of SMCs.^23^ The alteration of the blood flow associated with the mechanical injury induces SMC migration from the media towards the intima, and their proliferation. Combined with our *in vitro* results, which showed that S100A4 affected migration and not SMC proliferation or differentiation status, one can assume that, after 4 weeks of response to injury, an impaired migratory activity in S100A4 KO mice was masked by prominent proliferation, typical process of lesion formation after carotid artery ligation.^23^ Interestingly, in a similar model, 7 days after partial carotid artery ligation, single cell RNA sequencing showed that S100A4 was expressed at high levels in ECs before undergoing cell transitions and then was downregulated.^25^ Therefore, earlier time points might be worth analyzing, since S100A4 appears to play a role in the initial periods of the pathology.

One of the limitations of full KO models is the fact that these animals are completely devoid of the protein of interest. Compensatory mechanisms might have been activated, or other proteins might rise up to replace its functions. Different phenotypes may result from a hemizygous deletion of S100A4 and should be examined further.

S100A4 is commonly addressed as an inducer of migration in various types of cells, such as cancer cells,^26^ macrophages,^14^ ECs^27^ and SMCs.^28,29^ In our model, we have demonstrated that S100A4 KO SMCs exhibited a lower migratory capacity. As an intracellular protein, S100A4 is known to influence cell migration by binding various cytoskeletal proteins, such as non-muscle myosin-IIA and rhotekin,^15^ or modulating proteins involved in cell adhesion such as E-cadherin 5^30^ and liprin β1.^31^ Noticeably, both S100A4 KO and WT mice responded to PDGF-BB and PDGF-BB/rS100A4 to a similar extent. Moreover, rS100A4 did not influence SMC migration, contrasting with the work of Nagata et al.^7^, which showed that monomeric mouse rS100A4 enhanced the migratory activity of commercially available mouse aortic SMCs. This discrepancy could be due to differences in S100A4 protein conformation and/or SMC cultures.

Porcine coronary artery R-SMCs, typical of the synthetic phenotype, express S100A4 whereas cytoskeleton proteins, such as α-SMA, SMMHCs, smoothelin and desmin, are downregulated, and their proliferative activity increased when compared with S-SMCs, typical of the contractile phenotype.^3,4,28^ In this study, when characterizing the WT and S100A4 KO mouse SMC populations, no differences neither in morphology, in α-SMA and SMMHCs organization and expression level, nor proliferation were observed. On the other hand, no phenotypic switch was observed in S-SMCs treated with rS100A4, an effect only achieved when combining S100A4 with PDGF-BB.^6^ One possible explanation is that, although we detected a basal expression of S100A4 in WT SMCs, it might not be sufficient to elicit significant cytoskeleton and phenotypic changes in our model. Moreover, as it was not assessed whether S100A4 was being released into the extracellular medium of cultured WT SMCs, we cannot exclude that the S100A4-related effects (i.e. cell migration) are dependent on its release and action as an auto-paracrine molecule, as suggested by other groups.^32,33^ Even if S100A4 overexpression enhanced proliferation ^7^ or its downregulation inhibited cell growth,^34^ several groups refer to S100A4 as a motility stimulator rather than a mitogenic factor.^35–37^

Contrasting results have been published when defining the role of S100A4. For example, some cancer cell lines would respond to S100A4 by increasing proliferation while others did not.^38^ Moreover, extracellular S100A4 promotes the glycolytic pathway in a well-differentiated melanoma cell line (Melmet5), while it did not produce the same effect in a similar but poorly-differentiated Melmet1 cell line.^39^ It has been suggested that these differences are due to differential expression and pathway activation dependent on the differentiation status of the cell lines.^40^ Although the expression of SMMHCs in our primary SMCs might be an indicator of their differentiation status, we do not know how well differentiated they are in comparison with native medial SMCs. Notwithstanding, the above-mentioned studies were performed in tumor cell lines, which greatly differ from primary SMCs.

We treated S100A4 KO and WT SMCs with PDGF-BB, a known inducer of the SMC phenotypic switch,^2,41^ in order to understand whether the absence of intracellular S100A4 could affect some of the pathways responsible for this phenomenon. As expected, PDGF-BB, and to a greater extent, PDGF-BB plus rS100A4, promoted proliferation and downregulated α-SMA and SMMHCs expression. However, no difference was observed in both proliferation and SMC differentiation markers between WT and S100A4 KO SMCs. In contrast, we previously showed that PDGF-BB alone was not sufficient to promote the phenotypic switch of porcine coronary artery SMCs, only achieved when in combination with rS100A4.^6^ This difference could be species dependent.

NFκB activation was achieved with extracellular S100A4^6,27,28,32^ in both WT and S100A4 KO SMCs, indicating that S100A4 expression in the intracellular compartment is not required for extracellular S100A4 pathway activation. The fact that S100A4 KO SMCs were able to respond to PDGF-BB and rS100A4 at similar extents suggests that medial S100A4 KO SMCs are as sensitive as WT SMCs to growth factors and other chemical cues during IT development.

In conclusion, we propose that S100A4 plays a key role in the migratory capacity of murine SMCs. The carotid artery ligation gives rise to SMC-rich lesions with no major inflammatory components, a mechanism that is allegedly required for the mediated effect of S100A4.^6,11^ The absence of effect in IT development observed in S100A4 KO mice is likely due to a prevailing proliferation over migration process in this model.

## Acknowledgments

This study was supported by the Swiss National Science Foundation grant # 310030_185370/1 and by the Foundation Centre de Recherches Médicales Carlos and Elsie de Reuter.

We thank Alexandre Serigado, Anita Hiltbrunner and Aman Ahmed-Mohamed for excellent technical assistance, and Pascal Azar for helpful disscussions.

